## Supplemental Information for "Repeated activation of preoptic area-recipient neurons in posterior paraventricular nucleus mediates chronic heat-induced negative emotional valence and hyperarousal states"

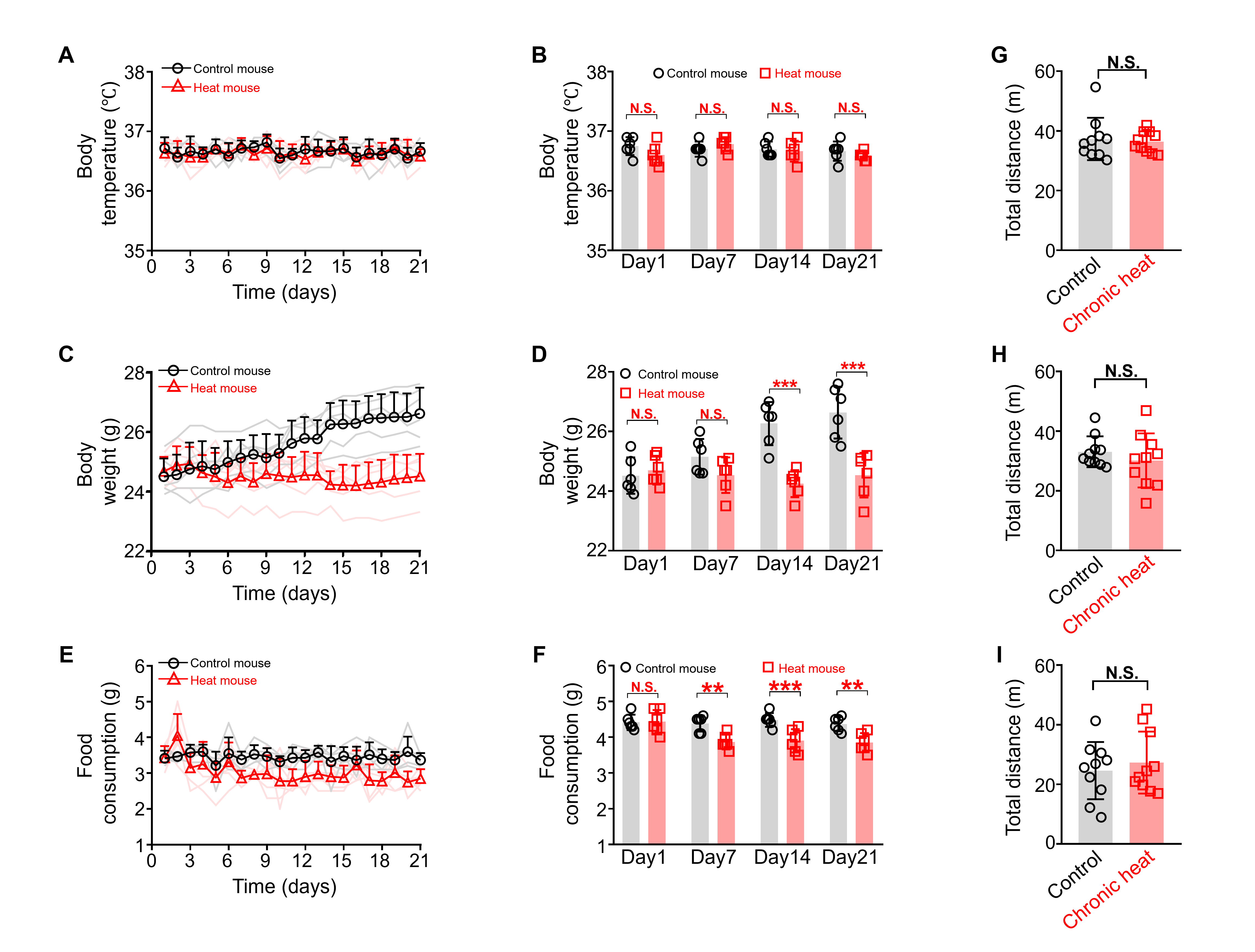

**Figure 1-figure supplement 1. The effect of chronic heat exposure on physiological states of mice and their motion activity during behavioral tests. (A, B)** Monitoring of body temperature of mice daily before heat exposure for both groups and statistical comparison (n=6 mice in each group, Two-way repeated measures ANOVA with Sidak post-hoc test; F(3, 40)=1.422, *p*=0.2505). **(C, D)** The effect of chronic heat exposure on mice’s body weight for both groups and statistical comparison (n=6 mice each, Two-way repeated measures ANOVA with Sidak post-hoc test; F(3, 30)=28.75, ****p*<0.001). **(E, F)** The effect of chronic heat exposure on mice’s food consumption for both groups and statistical comparison (n=6 mice each, Two-way repeated measures ANOVA with Sidak post-hoc test; F(3, 40)=3.781*,* **p=0.0177*). **(G-I)** Mice’s motion activity in the open field test (n=10 mice each, Mann-Whitney unpaired two-tailed U test; U=47*, p*=0.8534), the three-chamber test (n=10 mice each, Mann-Whitney unpaired two-tailed U test; U=41, *p*=0.5288), and the female encounter test (n=10 mice each, Mann-Whitney unpaired two-tailed U test; U=48, *p*=0.9118). **p*<0.05, ***p*<0.01, ****p*<0.001, N.S.: not important.

See also Supplementary Table S1 for further statistical information.

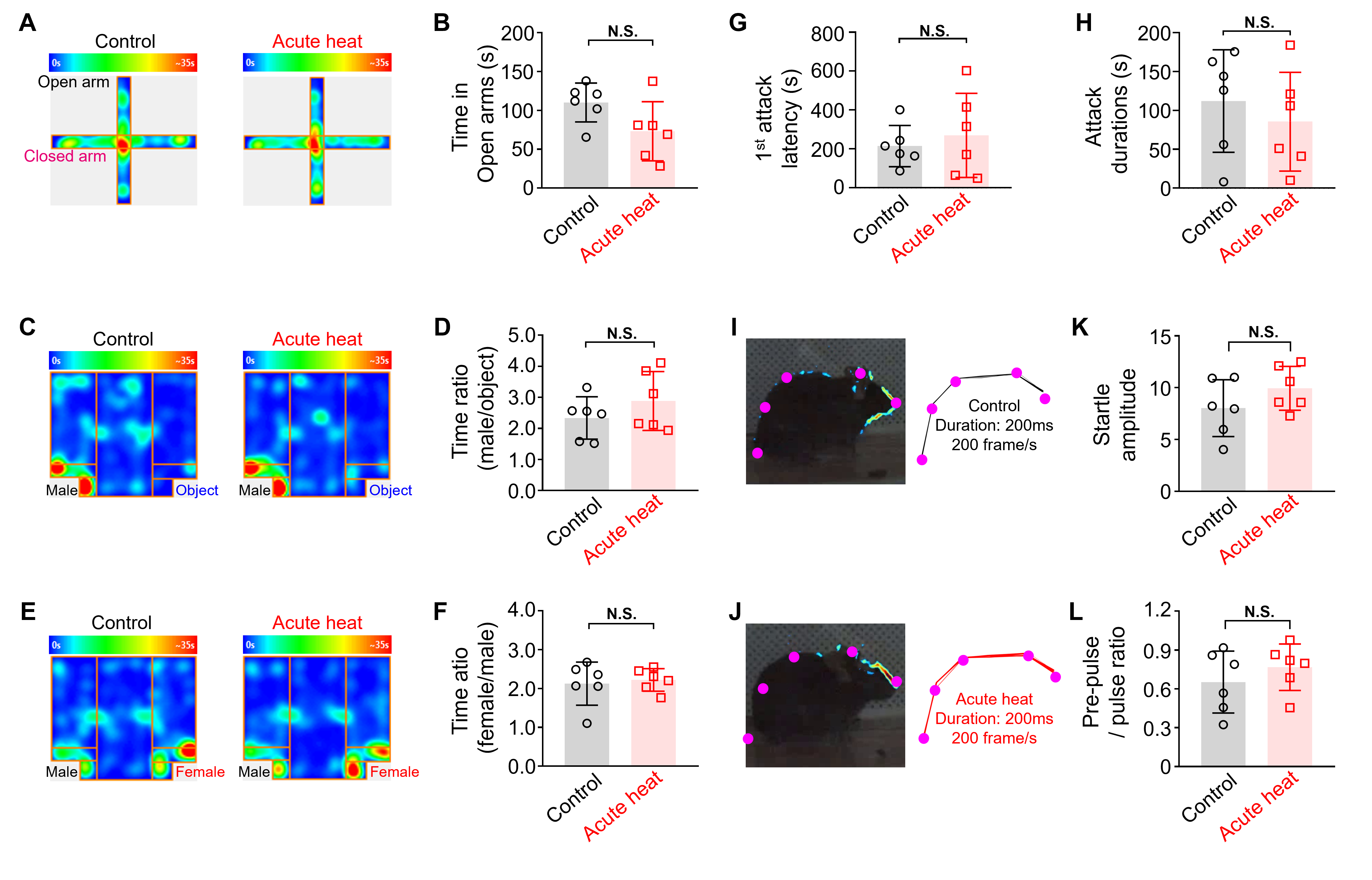
**Figure 1-figure supplement 2.** **Mice did not exhibit obvious changes of emotional valence and arousal states the day after acute heat exposure. (A, B)** The heatmap of representative tracking trace examples in EPM test and the time spent in the open arms (n=6 mice, Mann-Whitney unpaired two-tailed U test; U=8*, p*=0.132*)*. **(C, D)** The heatmap of representative tracking trace examples in TCT and the interaction time with an unfamiliar male mouse relative to the inanimate object (n=6 mice, Mann-Whitney unpaired two-tailed U test; U=13*, p*=0.4848*)*. **(E, F)** The heatmap of representative tracking trace examples in FET and the time surrounding the unfamiliar female mouse compared to an unfamiliar male mouse in FET compared for both groups (n=6 mice, Mann-Whitney unpaired two-tailed U test; U=17*, p*=0.9372*)*. **(G, H)** The 1^st^-time attack latency (n=6 mice, Mann-Whitney unpaired two-tailed U test; U=17*, p*=0.9372*)* and the attack durations (n=6 mice, Mann-Whitney unpaired two-tailed U test; U=13, *p*=0.4848) in the aggression test. **(I, J)** Visualized acoustic startle response examples (left panel) and corresponding labeled body parts’ skeletons (right panel) in ASR test from Control group and Acute heat group. **(K, L)** The startle amplitude (n=6 mice, Mann-Whitney unpaired two-tailed U test; U=10, *p*=0.2403) and the pre-pulse / pulse ratio (n=6 mice, Mann-Whitney unpaired two-tailed U test; U=13, *p*=0.4848) in ASR test. N.S.: not significant.

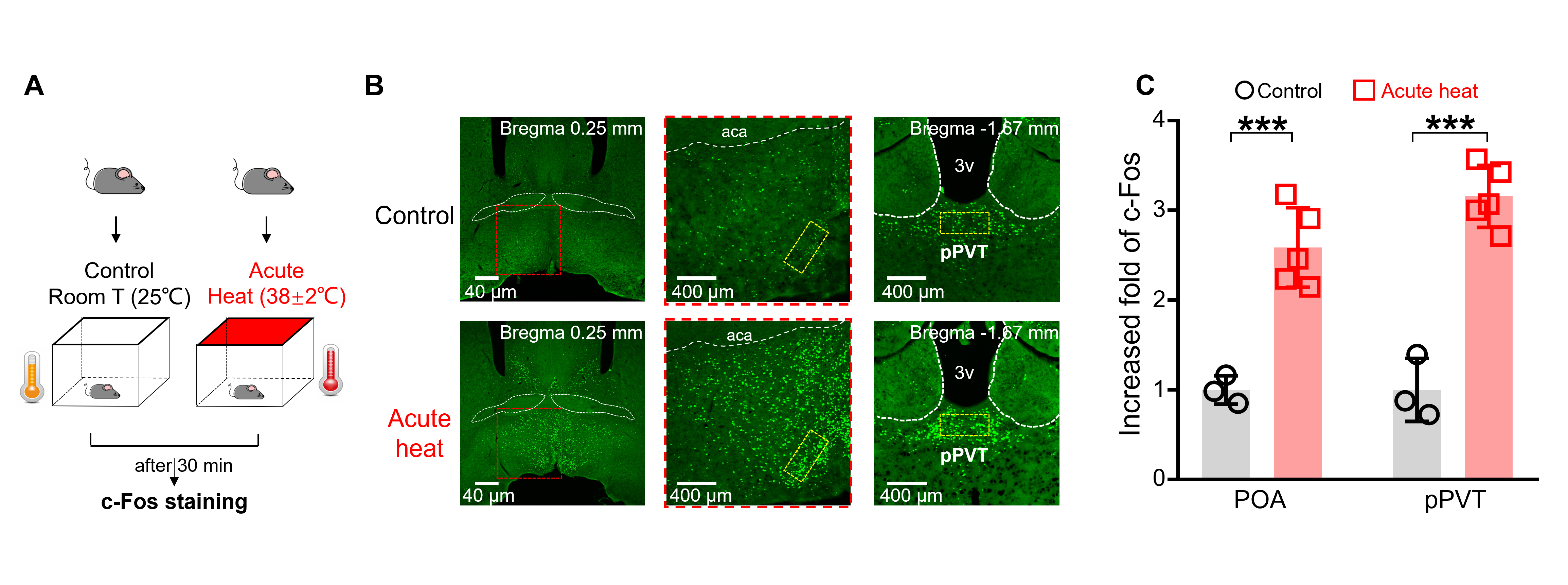
**Figure 2-figure supplement 1. pPVT was strongly activated after single-time heat exposure.** **(A)** Experimental schematics. Mice (n≥3 in each group) were randomly assigned to the Control and Acute heat groups, followed by Immunofluorescent staining and confocal imaging. **(B)** The representative microphotographs (each with a rectangular box (400 um [L] x 200 um [W]) for subsequent quantification) showed the c-Fos expression of the hypothalamic POA and pPVT for both groups. Magnification: left: 4x, middle: 10x, right: 10x. Scale bar: left: 40 μm, middle: 400 μm, right: 400 μm. **(C)** The c-Fos expression compared for both groups for both POA and pPVT (Two-way repeated measures ANOVA with Sidak post-hoc test; Interaction: F(1, 12)=2.103, *p*=0.2505; Heat treatment main effect: F(1, 12)=68.15*,* ****p*<0.001; Brain nuclei effect: F(1, 12)=5.159, *p*=0.0423; POA: Control vs. Acute heat: ****p*<0.001; pPVT: Control vs. Acute heat: ****p*=0.0008). ****p*<0.001.

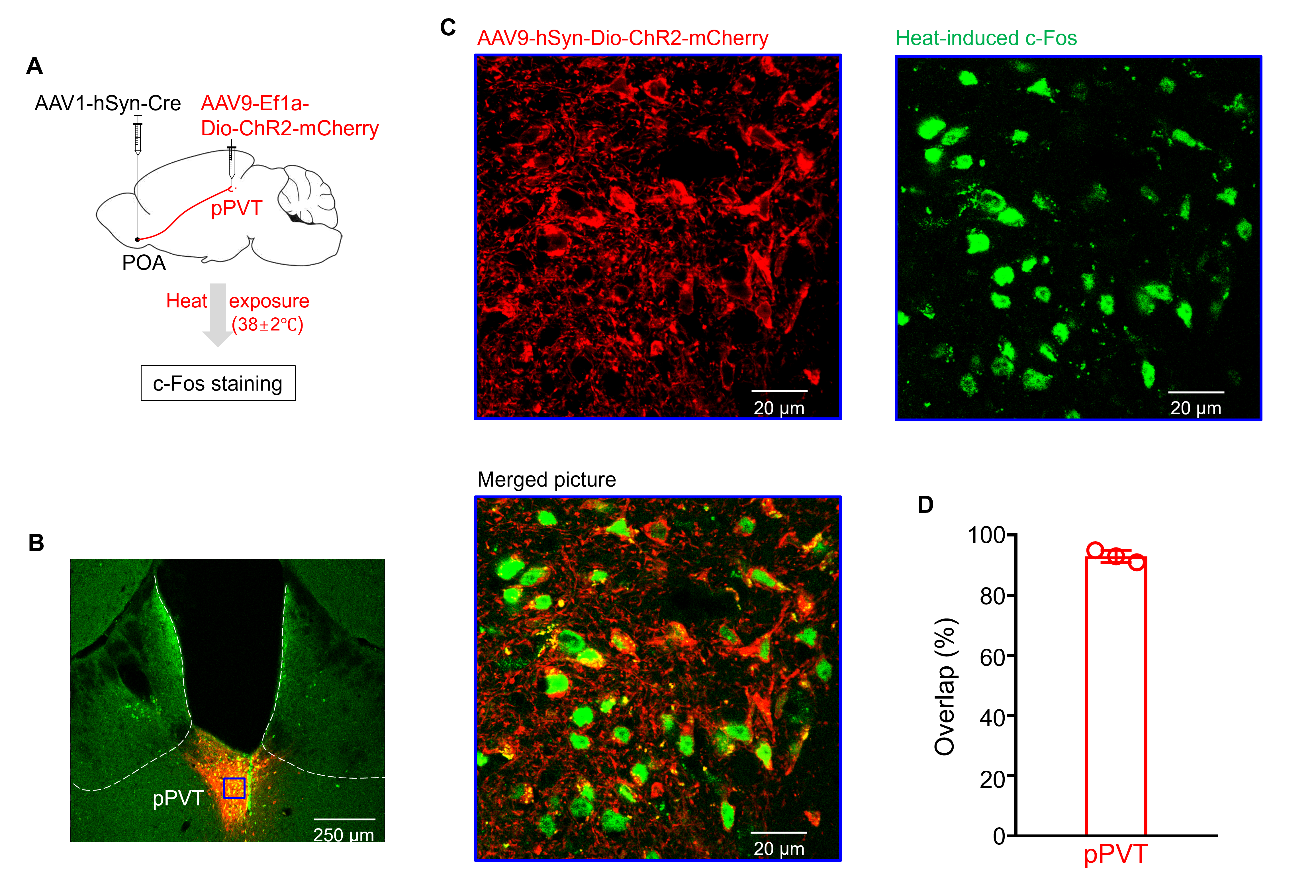
**Figure 2-figure supplement 2. POA recipient pPVT neurons were activated after heat exposure. (A)** Experimental schematics. Mice (n=3) were stereotaxically injected with AAV1-hSyn-Cre-EGFP in the POA and AAV9-Ef1a-Dio-ChR2-mCherry in the pPVT, followed by heat exposure and c-Fos staining. **(B)** The representative picture showed Cre-dependent ChR2-mCherry successfully expressed on pPVT neurons, embedded with c-Fos expression with green fluorescence. Magnification: 4x, scale bar: 250 μm. **(C)** The amplified pictures from the blue box in **(B)**, from left to right, respectively showed the expressions of ChR2-mCherry around the cell membrane, c-Fos with green fluorescence from the nuclei, and the merged picture of pPVT neurons. Magnification: 100x, scale bar: 20 μm. **(D)** The quantification of overlapping percentages of POA recipient pPVT neurons and heat exposure-induced c-Fos.

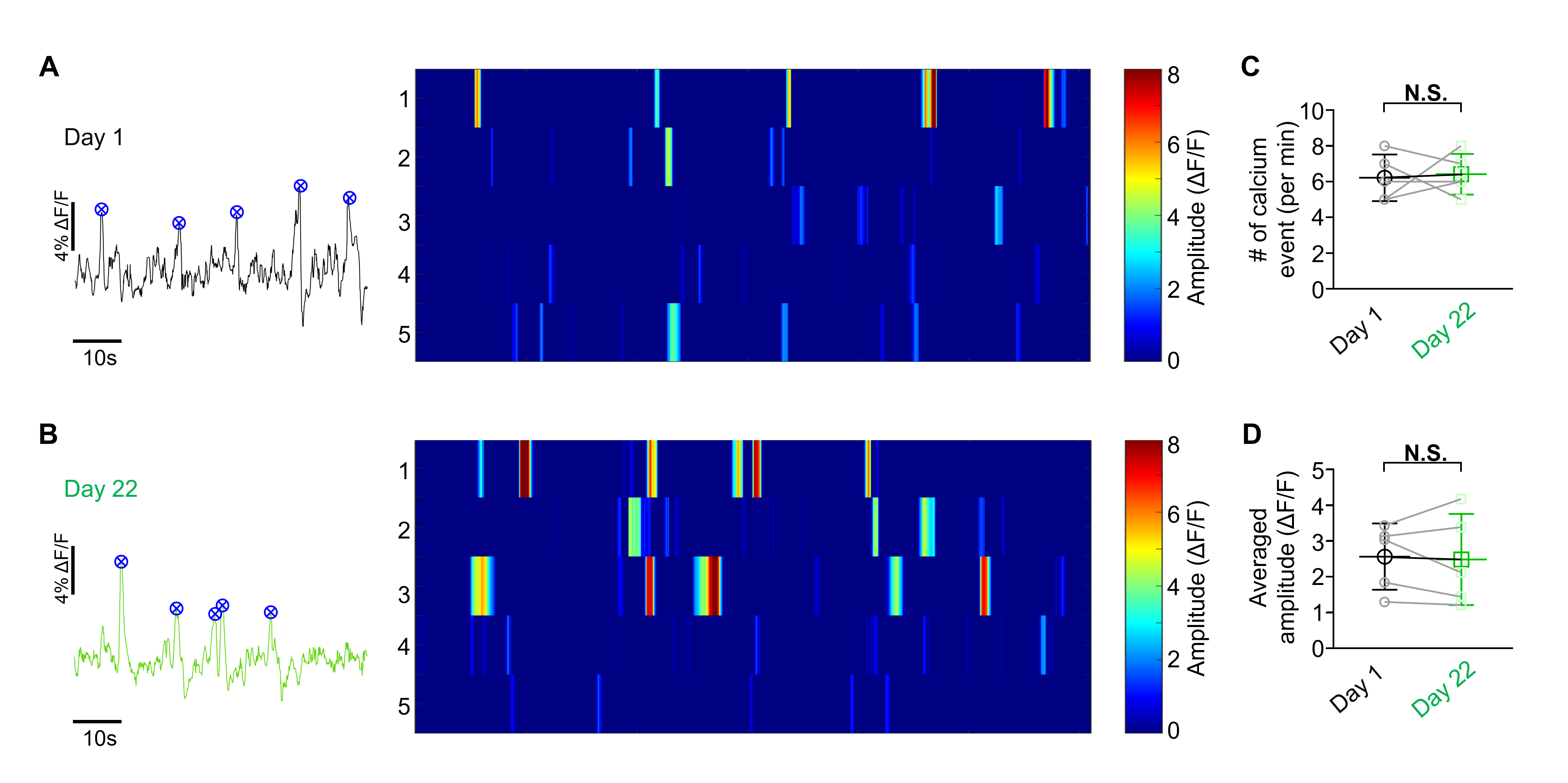
**Figure 3-figure supplement 1.** **The calcium activities of the POA recipient pPVT neurons were stable within our experimental period. (A, B)** The representative trace (left panel) and the overview of calcium events (right panel) of the Control group from day 1 and day 22. **(C)** The changes of frequency (Paired, parametric, two tailed t-test; t=0.2325, df=4, *p*=0.8276) and amplitude (Paired, parametric, two tailed t-test; t=0.2840, df=4, *p*=0.7905) of calcium events between day 1 and day 22. N.S.: not important.

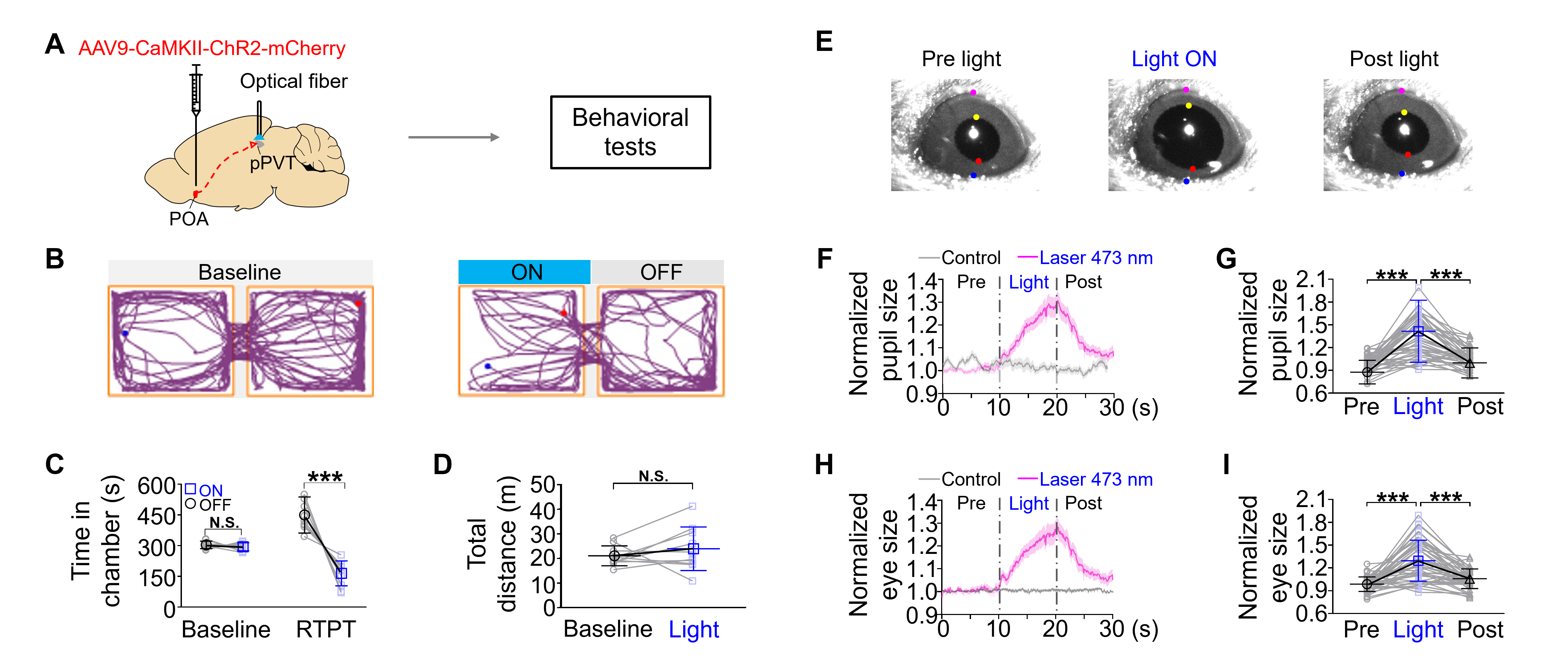
 **Figure 3-figure supplement 2.** **Optogenetic activation of POA excitatory neuronal terminals within pPVT produced aversive emotional valence and increased pupil size in mice. (A)** Experimental schematics. Mice (n=10) were stereotaxically injected with AAV9-CaMKII-ChR2-mCherry in the POA, followed by the implantation of optical fiber in the pPVT. **(B)** Representative pictures showed mice’s trajectory in the chamber without and with optogenetic activation, respectively. **(C)** The time of mice spent in the chamber associated with blue light stimulation (Two-way repeated measures ANOVA with Sidak post-hoc test; Interaction: F (1, 18)=33.59, ****p*<0.0001; Optical stimulation main effect: F(1, 18)=38.41, ****p*<0.0001; Chamber effect: F(1, 18)=1.42, *p*=0.2489; Baseline: OFF vs. ON: *p*=0.9513; Real-time place test: OFF vs. ON: ****p*<0.001). **(D)** The locomotion activity during the real-time place test (Paired, parametric, two tailed t-test; t=1.006, df=4, *p*=0.3406). **(E)** The representative pictures showed the pupil of mice pre-, during-, and post-light stimulation. **(F, G)** The changes of the pupil diameter (the vertical distance between yellow and red dots) of mice when the blue light was on and statistical comparison (One-way repeated measures ANOVA with Tukey post-hoc test; F(1.508, 67.88)=56.28, ****p*<0.001; Pre vs. Light: ****p*<0.001; Light vs. Post: ****p*<0.001). **(H)** The eye diameter (the vertical distance between purple and blue dots) of mice when the blue light was on and statistical comparison (One-way repeated measures ANOVA with Tukey post-hoc test; F(1.378, 62.01)=47.58, ****p*<0.001; Pre vs. Light: ****p*<0.001; Light vs. Post: ****p*<0.001). ****p*<0.001, N.S.: not significant.

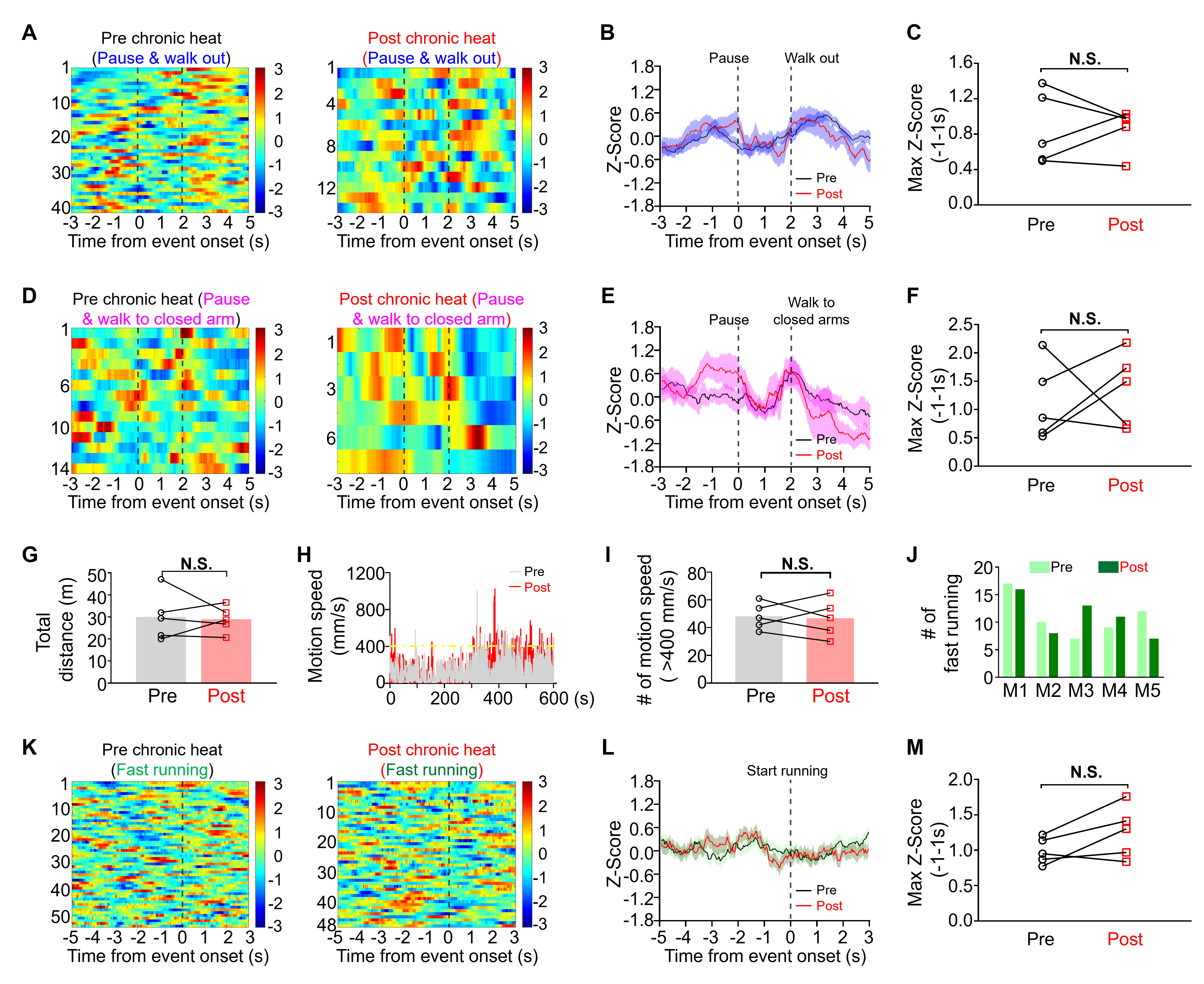
 **Figure 5-figure supplement 1.** **POA recipient pPVT neurons did not exhibit obvious changes in calcium activities when mice performed pause and walked to open arms, or walked to closed arms in the EPM, or displayed fast running in a heat-exposure-unrelated chamber.** **(A-C)** Heatmap showed the changes of calcium activities of the POA recipient pPVT neurons when mice performed pause and walked toward the open arms in the pre (n=41 trials from 5 mice) and post heat conditions (n=14 trials from 5 mice), Z-Score calcium signals, and statistical comparison (Paired, parametric, two tailed t-test; t=0.02394, df=4, **p*=0.98). **(D-F)** Heatmap showed the changes of calcium activities of POA recipient pPVT neurons when mice performed pause and walked to the closed arms in the pre heat (n=14 trials from 5 mice) and post heat conditions (n=7 trials from 5 mice), Z-Score calcium signals, and statistical comparison (Paired, parametric, two tailed t-test; t=0.5117, df=4, **p*=0.6358). **(G)** The changes in the total distance (Paired, parametric, two tailed t-test; t=0.2717, df=4, *p*=0.7993). **(H)** The changes of motion speed for pre and post conditions from one representative mouse. **(I)** The number of instances where motion speed exceeded 400 mm/s (Paired, parametric, two tailed t-test; t=0.2597, df=4, *p*=0.8079).**(J)** The number of fast running episodes compared for the pre and post heat conditions when mice were placed in a chamber unrelated to heat exposure. **(K-M)** Heatmap showed the calcium activities of the POA recipient pPVT neurons when mice performed fast running in the pre (n=54 trials from 5 mice) and post heat conditions (n=48 trials from 5 mice), Z-Score calcium signals, and statistical comparison (Paired, parametric, two tailed t-test; t=2.114, df=4, **p*=0.1021). N.S.: not important.

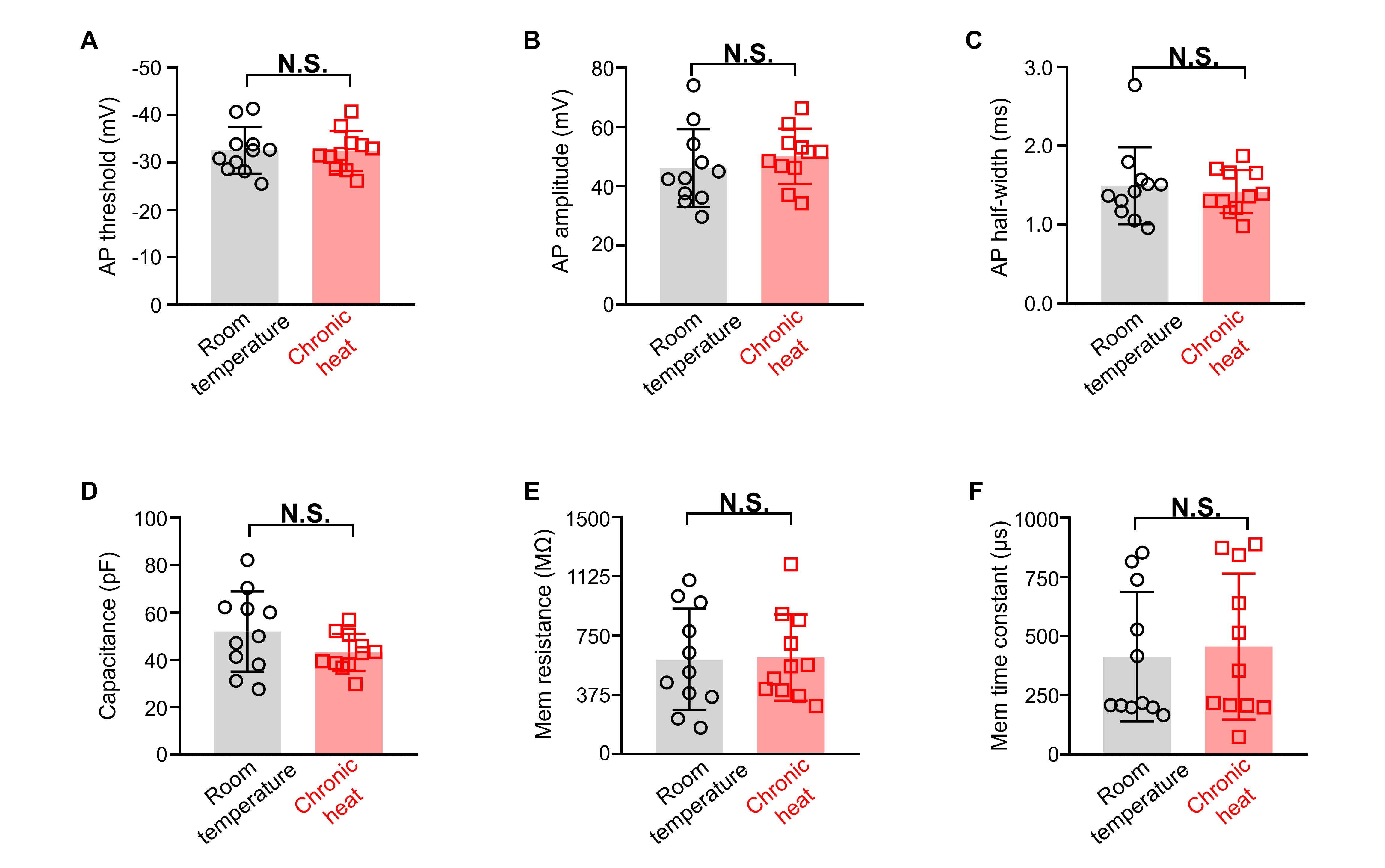
 **Figure 6-figure supplement 1. The effect of chronic heat exposure on intrinsic properties of pPVT neurons. (A-F)** The changes of half-width (Mann-Whitney unpaired two-tailed U test; U=58, *p*=0.8977), threshold (Mann-Whitney unpaired two-tailed U test; U=43, *p*=0.2703), peak amplitude (Mann-Whitney unpaired two-tailed U test; U=58, *p*=0.8977), capacitance (Mann-Whitney unpaired two-tailed U test; U=42, *p*=0.2426), membrane resistance (Mann-Whitney unpaired two-tailed U test; U=57, *p*=0.847) and membrane time constant (Mann-Whitney unpaired two-tailed U test; U=52.5, *p*=0.6177) of the action potential. N.S.: not significant.

**Figure 1-table supplement 1**

| **Fig. #** | **Statistical test** | **Factors, Degree of freedom & F/W value** | **p-value** | **Significance** |
| --- | --- | --- | --- | --- |
| **S1B** | Two-way repeated measures ANOVA with Sidak post-hoc test | Interaction: F (3, 40) = 1.422 | 0.2505 | N.S. |
|  |  | Day factor: F (3, 40) = 1.341 | 0.2747 | N.S. |
|  |  | Chronic heat treatment: F (1, 40) = 1.232 | 0.2736 | N.S. |
|  |  | Day 1: ctrl vs. heat | 0.2736 | N.S. |
|  |  | Day 7: ctrl vs. heat | 0.7849 | N.S. |
|  |  | Day 14: ctrl vs. heat | 0.9906 | N.S. |
|  |  | Day 21: ctrl vs. heat | 0.7849 | N.S. |
| **S1D** | Two-way repeated measures ANOVA with Sidak post-hoc test | Interaction: F (3, 30) = 28.75 | <0.001 | *** |
|  |  | Day factor: F (3, 40) = 5.376 | 0.0033 | ** |
|  |  | Chronic heat treatment: F (1, 40) = 36.99 | <0.001 | *** |
|  |  | Day 1: ctrl vs. heat | 0.9806 | N.S. |
|  |  | Day 7: ctrl vs. heat | 0.3643 | N.S. |
|  |  | Day 14: ctrl vs. heat | <0.001 | *** |
|  |  | Day 21: ctrl vs. heat | <0.001 | *** |
| **S1F** | Two-way repeated measures ANOVA with Sidak post-hoc test | Interaction: F (3, 40) = 3.781 | 0.0177 | * |
|  |  | Day factor: F (3, 40) = 4.135 | 0.0121 | * |
|  |  | Chronic heat treatment: F (1, 40) = 30.97 | <0.001 | *** |
|  |  | Day 1: ctrl vs. heat | >0.9999 | N.S. |
|  |  | Day 7: ctrl vs. heat | 0.0035 | ** |
|  |  | Day 14: ctrl vs. heat | 0.0009 | *** |
|  |  | Day 21: ctrl vs. heat | 0.0035 | ** |

**Figure 1-video supplement 1.** **Representative video showed the delivery of a 105 dB sound stimulus within 200 ms evoked an obvious body fluctuation in the chronic heat-exposed mouse.**

**Figure 3-video supplement 1. Representative video showed the enlargement of both the pupil and eye sizes of the head-fixed mouse during blue light stimulation of POA excitatory terminals within pPVT.**
